# Inferring TF activities and activity regulators from gene expression data with constraints from TF perturbation data

**DOI:** 10.1101/2020.05.25.108654

**Authors:** Cynthia Ma, Michael R. Brent

**Affiliations:** Center for Genome Sciences and Systems Biology, Washington University School of Medicine, St. Louis, MO 63110; Department of Computer Science and Engineering, Washington University, St. Louis, MO 63108; Department of Genetics, Washington University School of Medicine, St. Louis, MO 63110

**Keywords:** transcription factor activity inference, transcriptional regulatory networks, network component analysis (NCA), gene expression, TF perturbation, yeast

## Abstract

**Background:** The activity of a transcription factor (TF) in a sample of cells is the extent to which it is exerting its regulatory potential. Many methods of inferring TF activity from gene expression data have been described, but due to the lack of appropriate large-scale datasets, systematic and objective validation has not been possible until now.

**Results:** Using a new dataset, we systematically evaluate and optimize the approach to TF activity inference in which a gene expression matrix is factored into a condition-independent matrix of control strengths and a condition-dependent matrix of TF activity levels. These approaches require a TF network map, which specifies the target genes of each TF, as input. We evaluate different approaches to building the network map and deriving constraints on the matrices. We find that such constraints are essential for good performance. Constraints can be obtained from expression data in which the activities of individual TFs have been perturbed, and we find that such data are both necessary and sufficient for obtaining good performance. Remaining uncertainty about whether a TF activates or represses a target is a major source of error. To a considerable extent, control strengths inferred using expression data from one growth condition carry over to other conditions. As a result, the control strength matrices derived here can be used for other applications. Finally, we apply these methods to gain insight into the upstream factors that regulate the activities of four yeast TFs: Gcr2, Gln3, Gcn4, and Msn2. Evaluation code and data available at https://github.com/BrentLab/TFA-evaluation

**Conclusions:** When a high-quality network map, constraints, and perturbation-response data are available, inferring TF activity levels by factoring gene expression matrices is effective. Furthermore, it provides insight into regulators of TF activity.

## BACKGROUND

The activity level of a transcription factor (TF) in a given cell is the extent to which it is exerting its regulatory potential on its target genes. Cells process information, in part, by changing the activity levels of TFs, thereby changing the transcription rates of their target genes. Changes in TF activity can occur by several molecular mechanisms, including transcriptional and post-transcriptional regulation of the gene that encodes the TF, relocalization of the TF into or out of the nucleus, covalent modification of the TF such as phosphorylation, and non-covalent binding of the TF by other proteins. Because it has multiple molecular mechanisms, TF activity is difficult to measure directly. However, it may be possible to infer changes in TF activity level from changes in the expression levels of the TF’s target genes (1-25). If successful, this would provide insights into which TFs are involved in the transcriptional response to a given stimulus, such as a drug, extracellular signal, or nutrient influx. In principle, TF activity inference could also be used to predict the transcriptomic effects of direct perturbations of TF activity levels, such as deletion or over-expression. Finally, inferred activity levels could be used to improve TF network mapping (5, 12, 14, 16, 22-24, 26-33).

Most algorithms for inferring TF activity (TFA) from gene expression data fit the parameters of a mathematical model to the expression data (1, 5-8, 10, 12-14, 17-19, 21, 23, 24, 29, 31). These models include parameters representing TFA levels, the values of which vary from one biological sample to another. Some models also include parameters that are constant across samples but vary as a function of the TF and target gene. These parameters reflect factors such as the affinity of the TF for sites in the promoter of each gene. In some approaches, these parameters, called *control strengths* (CSs), are obtained directly from TF binding data (12, 17, 19) or by scanning models of TF binding specificity across promoters (7, 31, 34). In other approaches, they are treated as unknowns and obtained by fitting the model to gene expression data (1, 5, 6, 8, 14, 18, 24). In this case, gene expression is typically modeled as depending linearly on the TFA levels and on the CSs (1, 8, 14, 24), so the model as a whole is bilinear. More highly parameterized, non-linear models that more closely reflect the underlying biochemistry have also been tried when modeling a small number of TFs (5, 6, 18). Because it has relatively few parameters and can be fit by a simple algorithm, we focus on the bilinear framework.

To infer the activity of a set of TFs from the expression of their target genes, an inference algorithm must know at least some of the targets of each TF. We refer to this input as a *TF network map* (35). TF network maps link each TF to the targets it has the potential to regulate directly, given the right conditions. These maps are qualitative, so they can be represented by binary adjacency matrices. Fitting the bilinear model yields a control strength for each edge of the input map. Multiplying these CSs by the TFA levels inferred for a sample of cells yields a sample-specific network map showing how strongly each TF is influencing the expression of each of its targets in that sample. In previous work featuring inferred CS values, qualitative network maps have mostly been constructed from binding location data obtained by chromatin immunoprecipitation (ChIP) (1, 5, 6, 8, 10, 20-24, 28, 32, 33, 36, 37). Garcia-Alonso et al. reported that a manually curated network map performed best for TFA inference in human (38), but curated networks include very few TFs and are not available for most organisms. Here, we analyze the effects on TFA inference accuracy when comparably-sized networks are constructed from various high-throughput data sources.

In most previous studies, inferred TFA values were allowed to be positive or negative and their absolute value was interpreted as the magnitude of activity change relative to some reference sample. Thus, a smaller absolute TFA did not indicate less activity, but rather less change relative to the reference. As a result, the TFA levels did not distinguish between increasing and decreasing activity. Furthermore, the signs of the CS values had no meaning. Here, we propose, evaluate, and optimize a version of the bilinear approach in which TFA values are constrained to be non-negative, so that zero represents no activity, equivalent to deletion of the gene encoding the TF. We also include parameters representing the expression of each when all its regulators have activity zero (*baselines*). The combination of baseline expression levels with the non-negativity constraint on TFA values differentiates our model from previously proposed models. Together, they make the parameters interpretable. Positive control strength indicates that the TF activates the target and negative control strength indicates that it represses the target. If a TF’s activity is larger in one sample than in another, then the TF is more active in the former sample than in the latter. We make extensive use of gene expression data after direct perturbations of TF activities (23, 28, 39), constraining each control strength parameter to be positive (activating) or negative (repressing) based on the direction in which the target gene’s mRNA level changes when the TF is perturbed. If the gene encoding a TF is deleted in a sample, the TF’s activity is held at zero; if it is overexpressed, the TF’s activity is constrained to be greater than its activity in unperturbed samples

Evaluating the effects of various mathematical models, network mapping procedures, and perturbation-derived constraints, requires accuracy metrics that are objective, quantitative, and available for large numbers of TFs. This poses a challenge because TFA cannot be directly measured. As a result, most attempts to validate TFA inference algorithms have been small-scale and often qualitative, highlighting successes with just a few TFs. Some validation efforts have been based on inferring significant differential activity in a handful of samples subjected to stressors (2) or small molecules known to affect a particular TF’s activity (3, 20, 40). Others have been based on inferring activity patterns that appear to match the periodicity of cell cycles (1, 5, 6) or using changes in the nuclear localization of a GFP-tagged TF as a proxy for TFA (2). Other evaluation efforts have been based on internal consistency measures (41), TF activity perturbations (2, 38, 42), or identification of TFs important for proliferation of cancer cells (3, 4, 7, 9-11, 13, 15, 17, 19, 20, 25, 40, 42, 43). In this work, we take advantage of two independent, high-quality perturbation datasets in *Saccharomyces cerevisiae* to present multiple quantitative, large-scale validation metrics. By using one dataset for network construction and constraint generation, and the other for validation, we reveal which high-throughput data types are most valuable for TFA inference in the matrix factorization framework, identify best practices for achieving high accuracy, and show that given the right input data, TFA inference works reasonably well.

## RESULTS

### Mathematical model of gene expression

We use a simple model in which the log expression level of a gene in a given sample is determined by its baseline expression level, when none of its regulators are active, plus the sum of the influences of all the TFs that regulate it. The influence of each TF is a product of the strength with which the TF regulates that gene (*control strength*) and the TF’s activity in that sample:

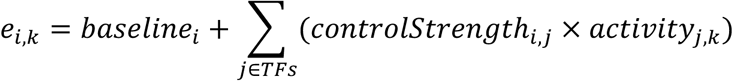

where *e*_*i,k*_ is the log expression level of gene *i* in sample *k, baseline*_*i*_ is the expression level of gene *i* absent any influence from TFs, *controlStrength*_*i,j*_ is the condition-independent potential of TF *j* to activate or repress gene *i*, and *activity*_*j,k*_ is the activity level of TF *j* in sample *k*. In matrix notation, **E** = **CS** • **TFA**, where **E** is a gene expression matrix (genes by samples), **CS** is a matrix of control strengths (genes by TFs) augmented to incorporate baselines, **TFA** is a matrix of TF activity levels (TFs by samples), and • indicates matrix multiplication (Fig. 1). Fitting the CS and TFA matrices to expression data is equivalent to factoring the expression matrix, under the constraints that CS signs are predetermined, TFA is non-negative, and the activities of perturbed TFs are constrained according to the perturbation. The model is fit by minimizing the sum of squared errors with this equation. This can be done by alternating linear regression (1, 6, 19, 24, 39) or with a more general nonlinear optimization algorithm (44). See Methods for details.

**Figure 1.**
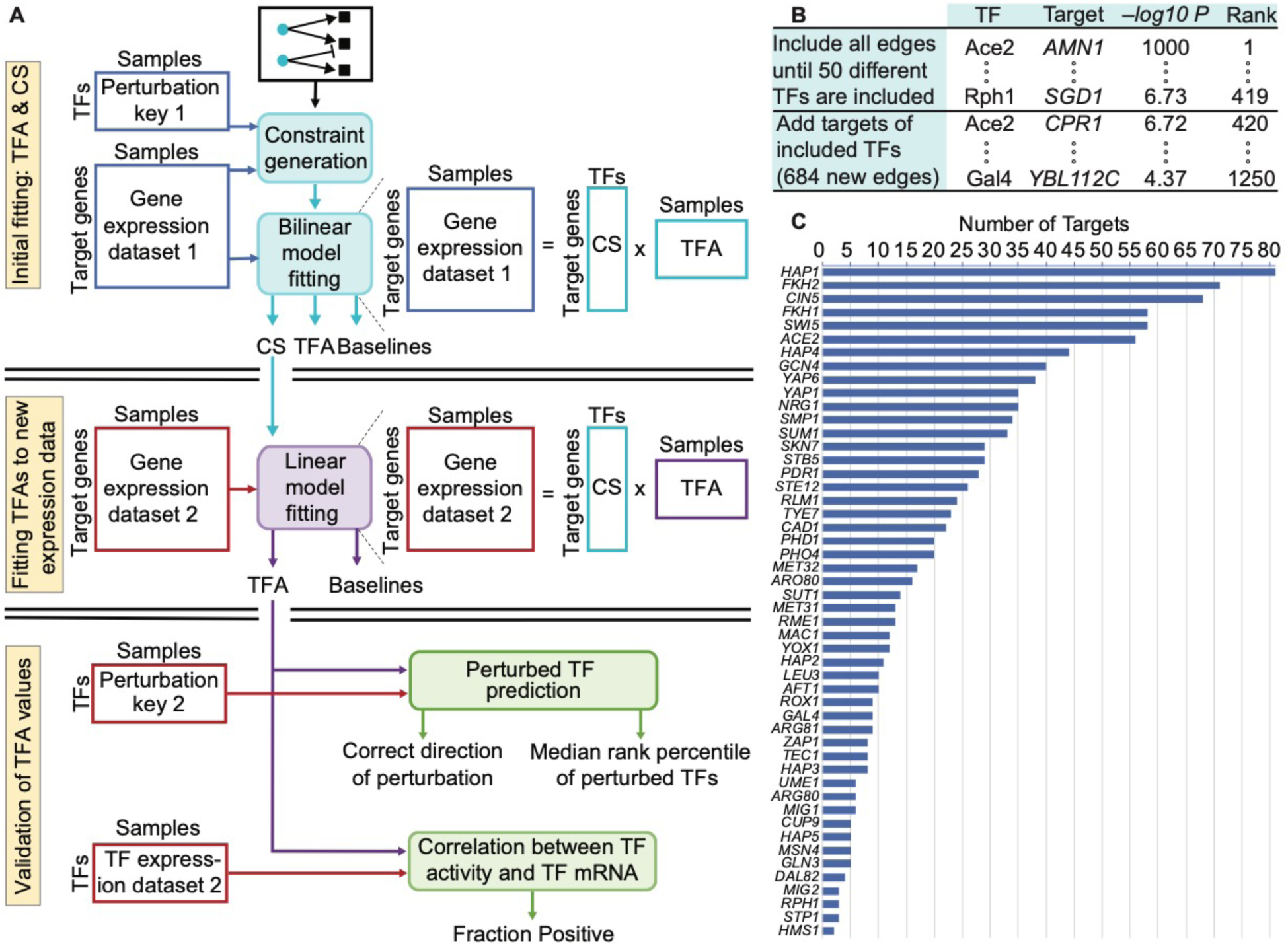
Evaluation framework and ChIP-based network construction. A. Overview of three-stage model fitting and TFA evaluation procedure. Gene expression levels and the perturbation key from dataset 1 are used only in the initial fitting. The CSs inferred in the initial fitting are fixed while the TFAs and baselines are refit to the target gene expression levels from dataset 2. The mRNA levels of the TFs and the perturbation key from dataset 2 are used only for evaluation. B. Illustration of how edges were selected for the ChIP-based network. All edges were ranked according to their -log *P*-value for the TF binding in the promoter of the target. Edges were selected in rank order until there was at least one edge from 50 different TFs. Lower-ranked edges were then selected for those TFs until rank 1250. After initial model construction, we removed any TFs with a single target and any set of TFs with identical target sets, along with their target genes. We then returned to the list and iteratively added edges that had previously been passed over until the network stabilized at 50 TFs. This yielded a network with 1,104 edges. C. The number of targets for each of the 50 different TFs in the ChIP network.

### Core evaluation metrics

Our approach to TFA evaluation relies on two independent expression data sets in which the activity of each TF has been individually perturbed. The first data set consists of expression profiles of strains in which a single TF was deleted from the genome (*TFKO* data) (45). The second consists of expression profiles collected 15 minutes after overexpression of a single TF was induced using the ZEV system (*ZEV* data) (46). These two data sets represent two very different growth conditions: TFKO profiles cells growing on synthetic complete medium in shake-flasks with no nutrient limitation, while ZEV profiles cells growing on minimal medium in phosphate-limited, continuous-flow chemostats. Two of our core evaluation metrics focus on whether TFA inference can determine which TF was perturbed in each sample and whether it was knocked out or induced. We refer to this information as the *perturbation key*. First, we use both the perturbation key and the expression profiles from one data set for network construction, constraint generation, and fitting (Fig. 1A, top). Next, we hold the control strengths from the initial fit fixed and refit the TFA and baseline parameters to the expression profiles in a second perturbation dataset (Fig. 1A, middle). Crucially, the perturbation key for the second dataset is not used for either fit, so it can be used as independent evaluation data (Fig. 1A, bottom).

We use three core evaluation metrics for TFA accuracy. The *direction of perturbation* metric is the fraction of samples in which the direction of perturbation (deletion or overexpression) is inferred correctly, given the identity of the perturbed TF. For the second metric, all TFs in a perturbation sample are ranked, starting from the one whose inferred activity increased most, relative to an unperturbed sample (for overexpression) or decreased most (for deletion). We expect the perturbed TF to be highly ranked. The median rank percentile, across all perturbation samples, is the *median rank percentile* metric. The *positive correlation* metric is the fraction of TFs with positive correlations between their activity levels and their mRNA levels. To ensure that TFA-mRNA correlations were not driven by a few outliers, we selected one thousand bootstrap sets of samples, calculated the TFA-mRNA correlation for each TF using only those samples, and reported the fraction of correlations that were positive, for each TF. We used the median fraction of positive correlations, across all TFs in a network, as the third metric. We do not expect accurate TFA values to yield 100% for this metric, since post-transcriptional regulation is a key determinant of TFA levels, but we do expect the fraction of positive correlations to be substantially greater than half.

The metrics reported below are averages after training on each dataset and testing on the other, with P-values combined by using Fisher’s combined probability test (47). The first fitting only used samples in which one of the network TFs was directly perturbed, while the second fitting also used samples in which other TFs were perturbed. For all analyses, we considered only the 179 TFs that were perturbed in both ZEV and TFKO datasets. For *median rank percentile*, inferred activity levels for each TF were log-ctransformed and then standardized across samples, converting them to Z-scores. This makes the activity levels of different TFs comparable.

### Constructing a ChIP-based network

Suppose binding location data are available for many of the TFs in a given organism, along with gene expression profiles from many samples. For now, assume that the expression data does not contain direct TF perturbations or, if it does, that the perturbation key is not used. To generate a network map as input for TFA inference, all possible TF-target *edges* can be ranked according to the strength of evidence that the TF binds in the promoter of the gene. We did this for yeast, ranking edges according to their negative log *P*-value in a comprehensive ChIP-chip dataset (48). This produced a single, global ranking of all edges involving all TFs (Fig. 1B). We then constructed a network as described in Figure 1B. All networks are described further in Supplementary Methods and are provided as supplemental files.

### Evaluating TFAs inferred from the ChIP network with correlation-based constraints

First, we evaluated the ChIP network without any constraints on the parameters (except non-negative TFA) and found that it did not perform better than chance on any metric (Fig. S1). Next, we tried adding constraints on the signs of the control strength parameters, based on the intuition that a positive correlation between the mRNA levels of a TF and a target suggests activation while a negative correlation suggests repression. We constrained the sign of each control strength to match the sign of that correlation (even if the correlation was not significant) using the dataset reserved for the initial fitting, which improved performance. We refer to the ChIP-based network with correlation-based constraints as ChIP-CC (File S1-4). Using ChIP-CC, the direction of activity change between a TF’s perturbation sample and the unperturbed sample was predicted correctly for 66% of TFs (*P* < 0.01, binomial test; Fig. 2A, left), the median rank percentile was 76.5% (P < 0.0001, binomial test; Fig. 2A, middle), and at least 65% of TFs’ activity levels were positively correlated with their mRNA levels in half the sets of bootstrapped samples (Fig. 2A, right; P < 0.01).

**Figure 2.**
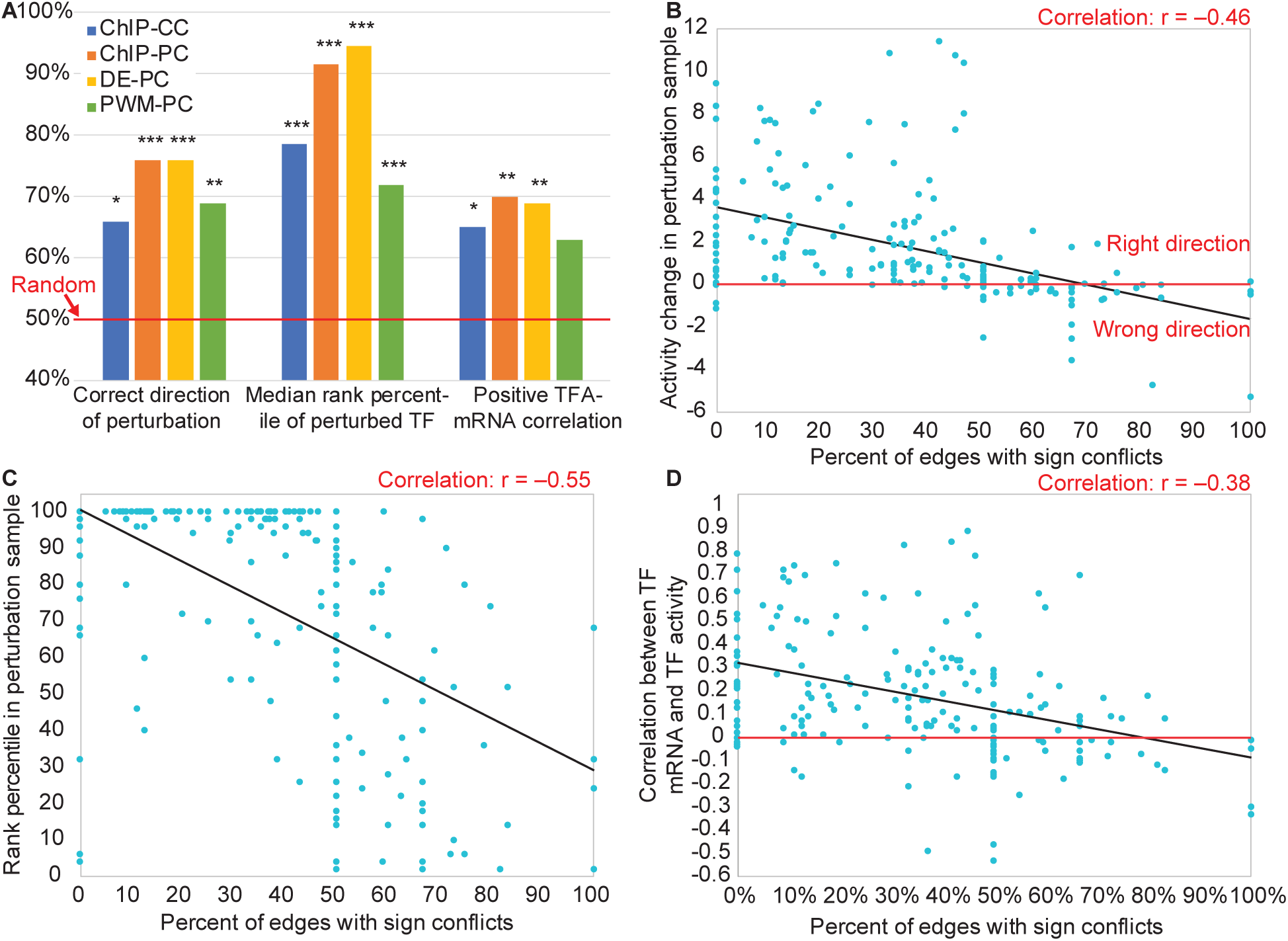
Determinants of TFA accuracy. A. Effects of network construction and constraint generation on TFA accuracy. Blue: ChIP network with correlation-based constraints. Orange: ChIP network with perturbation-based constraints. Yellow: Differential expression network with perturbation-based constraints. Green: Binding-specificity (PWM) network with perturbation-based constraints. Asterisks above the bars indicate magnitude of significance compared to a random model, with 1, 2, or 3 asterisks representing p-value thresholds of 0.01, 0.001, or 0.0001 B. Vertical axis: The activity of each TF in the sample in which it was perturbed minus its activity in the unperturbed sample, oriented so that higher is better. TFs plotted below the horizontal axis have been inferred to change activity in the wrong direction. Horizontal axis: The fraction of each TF’s targets for which the TFKO and ZEV data sets suggest conflicting CS signs. TFs with < 50% conflict edges are almost all predicted in the correct direction, while most TFs with > 50% conflict edges are not. C: Vertical axis: Rank percentile of the perturbed TF’s activity change in each perturbation sample (higher is better). Horizontal axis: Same as B. TFs with a higher percentage of conflict edges tend to be ranked lower. D: Vertical axis: median fraction of bootstrap samples in which a TF’s mRNA level and its inferred activity level are positively correlated (see main text). TFs with a higher percentage of conflicting edges tend to have low or negative correlation. B-D: Results from the 50-TF ChIP-PC and DE-PC networks in both train-test directions have been combined, but each individual set of 50 points showed similar, highly significant correlations.

### Adding constraints obtained from TF perturbation data

When expression data after direct TF perturbation are available, the perturbation key can be used to constrain both the CS signs and the activity of the TF perturbed in each sample, as described above. We evaluated the effect of using the perturbation key from the first data set to constrain the control strengths and activity levels during the first fit (Fig. 1A, top). The perturbation key for the second dataset is only used for evaluation (Fig. 1A, bottom), not during the second fit (Fig. 1A, middle). We call the ChIP network with perturbation-constraints ChIP-PC (File S7-10). Performance on all three metrics increased, relative to using correlation-based constraints (Fig. 2A, blue and orange bars).

### Generating network and constraints from TF perturbation data, without binding data

If expression data from direct TF perturbations is available, it is possible to build a network from the perturbation data rather than ChIP data. The same network building procedure is used, but instead of TF-target interactions being ranked by the strength of ChIP evidence, they are ranked by the absolute value of the log fold change of the target when the TF is perturbed (DE-PC, Files S13-16). The performance of this network with perturbation-based constraints was similar to that of ChIP-PC, with the biggest change being an increase in median rank percentile from 92% to 96% (Fig. 2A, red and orange bars). We conclude that differential expression data from direct TF perturbations are necessary and sufficient for accurate TFA inference performance -- binding location data are not necessary.

### Using binding specificity models in network generation

A popular source of data for building gene regulatory networks is models of TF binding specificity, typically represented as position weight matrices (PWMs) (2, 11, 15, 27, 31, 36). To test this approach, we ranked all possible TF-target interactions by the maximum, across all positions in the target gene’s promoter, of the negative-log P-value for presence of the TF’s motif, as defined in the ScerTF database (49) and scored by FIMO (50) (see Supplement). We then built a network from this ranking just as we did with ChIP-chip data and the differential expression data (Fig. 1B). Using this network, we optimized with perturbation-based constraints (PWM-PC, Files S19-22). This network performed significantly worse than both ChIP-PC and DE-PC networks (Fig. 2A, green bars).

### Increasing the number of TFs

To infer activity for more than 50 TFs, the input network maps can be extended by considering more than just the 1250 top ranked edges. To quantify the loss in performance from using lower ranked edges, we used blocks of 2000 edges of decreasing rank to make independent networks of 50 TFs (Supplemental Methods). Both ChIP-PC and DE-PC lost accuracy steadily as lower ranked edges were used. DE-PC lost accuracy more slowly in the first two metrics (Fig. 3A, B) while both networks lost ground at the same rate for TFA-mRNA correlation (Fig. 3C). Thus, DE-PC appears to be the better choice for building larger networks with more TFs.

**Figure 3.**
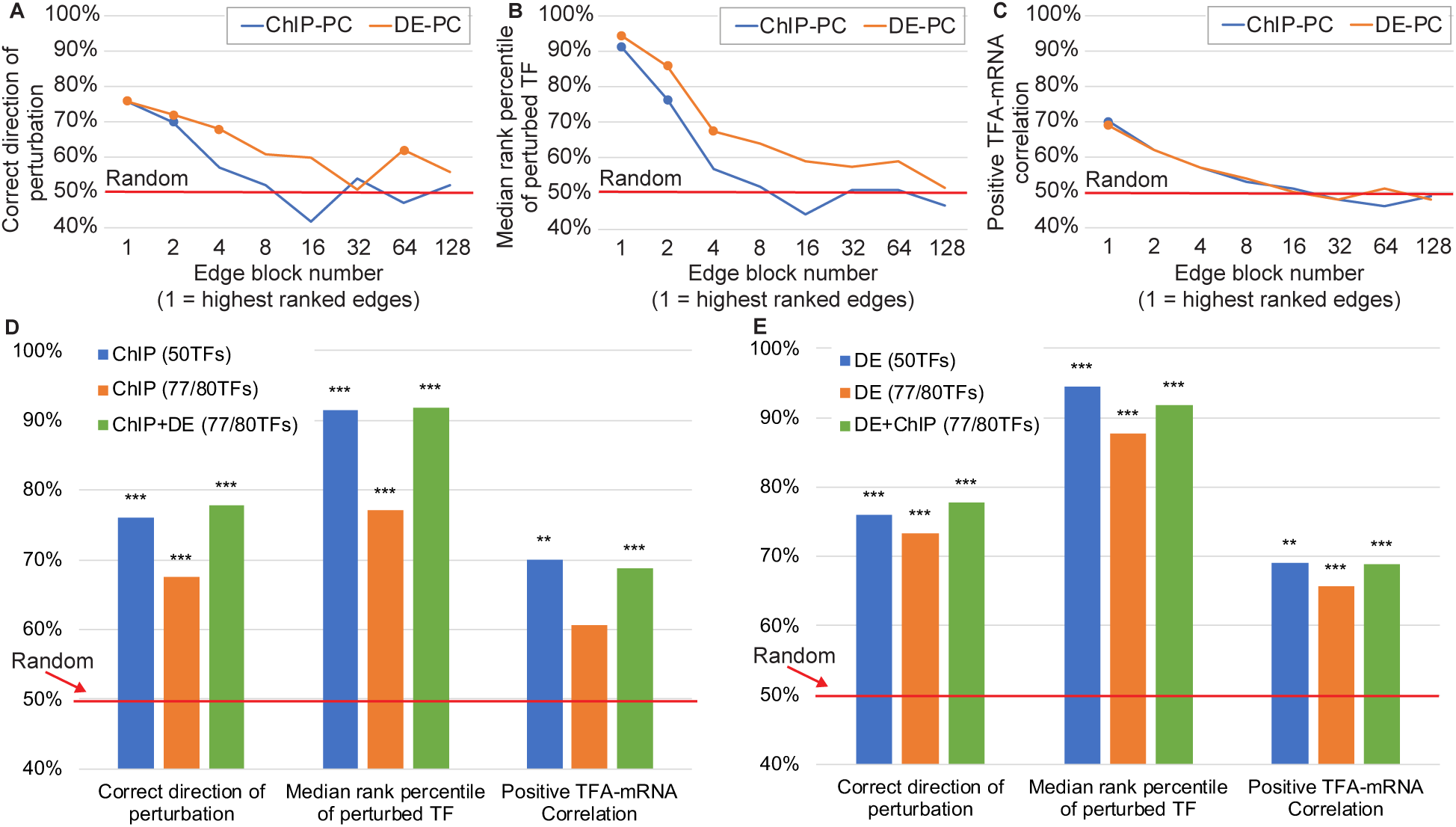
Effects of increasing the number of network TFs on accuracy. A-C: Accuracy metrics for networks constructed from the ChIP or DE edge lists by taking successively lower ranked edges. Edges were divided into blocks of 2,000 and blocks are plotted in an exponential series. For example, Block 1 is edges ranked 1-2,000 and Block 4 is edges ranked 6,001-8,000. Points are plotted for results that are significantly better than random (P < 0.001) A: Percent of TFs whose direction of perturbation is predicted correctly. B: Median rank percentile of the perturbed TF. C: Percent of TFs with a positive TF-mRNA correlation. In A and B, the ChIP-PC performance starts out similar to DE-PC, but it drops faster to no better than random in any measure by Block 4. D. Comparison of two ways of increasing the number of TFs in the network -- going further down the list of ChIP edges or using 50-TF ChIP and DE networks and averaging standardized TFAs of TFs that are in both networks. Consistent with panels A-C, performance degrades when lower ranked edges are included in the ChIP network. Inferring TFAs separately and averaging them, by contrast, yields performance on a larger network that is as good as performance on the smaller, 50-TF networks. E. Same as D, but blue and orange bars are for DE networks.

If both ChIP and TF perturbation data are available, another option is to infer activities separately for 50-TF ChIP-PC and DE-PC networks and combine the results, averaging the standardized activities for TFs that are in both networks. This provides activities for a larger number of TFs with accuracy that is better than including lower-ranked edges from a single data source (Fig. 3D, E, orange vs. green bars).

### Effect of optimizing control strengths on TFA accuracy

To determine how much optimizing control strengths contributes to the accuracy of inferred TFAs, we compared TFAs obtained by optimization of both TFA and CS matrices to TFAs obtained by using fixed control strengths of +1 for activation or -1 for repression (*signed binary* CSs). Signed binary CSs were also used in (10, 16, 23, 28). The optimized CSs performed slightly better on some metrics and some networks, but there was little difference overall (Fig. 4A, B). To investigate the potential value of CS optimization further, we considered two additional metrics.

**Figure 4.**
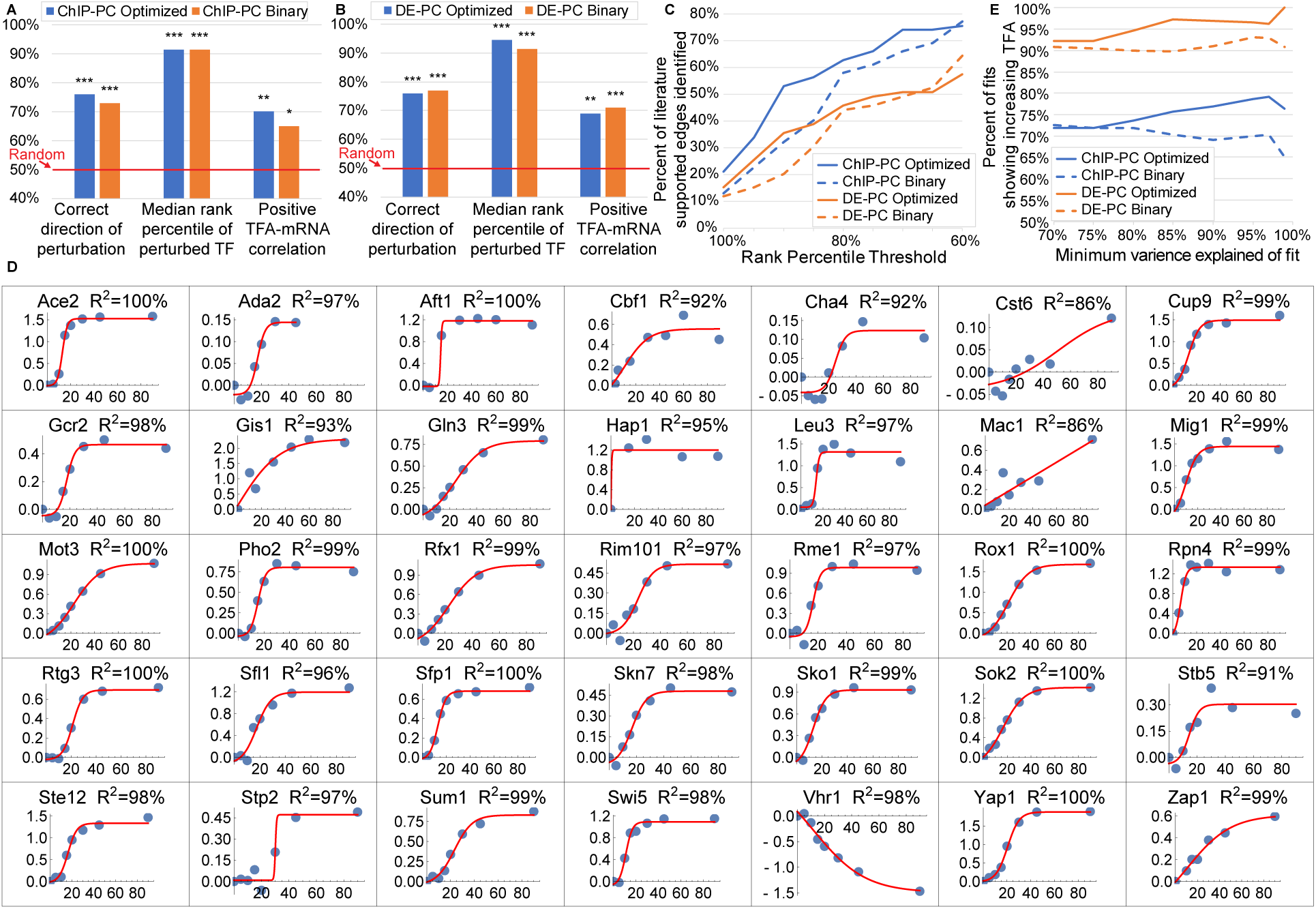
Impact of using a CS matrix optimized on a different data set versus using a signed binary CS matrix. A. ChIP-PC network. B. DE-PC network. C. Percent identification of literature-supported edges between TFA regulators and TFs, as a function of minimum rank percentile for identification. Solid lines: CS matrix optimized on the ZEV dataset and used to infer TFA’s in the samples in which a TF regulator was deleted. Dashed lines: Signed binary CS matrix. For TFs whose change in standardized log activity from WT ranks above 85th percentile, more literature supported edges are identified by using optimized CS matrices than by using signed binary matrices. D. Sigmoidal fits to log2 fold change of TFAs inferred for the ZEV time course data, using the DE-PC network and a CS matrix optimized on the TFKO dataset, relative to the 0min timepoint. Only fits with variance explained above 85% are shown. In all but one of the 35 fits, TF activity is correctly inferred to be increasing (97%). Only Vhr1 activity is inferred to change in the wrong direction, probably because 9 of its 11 targets have sign conflict (80%, see Fig. 2B-D). E. After fitting sigmoidal curves as in D and imposing various thresholds on the variance explained by the fit, the percentage of fits that correctly show increasing activity. The DE-PC network (orange lines) performs better than the ChIP-PC network (blue lines). For each network, using a CS matrix optimized on the TFKO data (solid lines) generally shows better performance than using a signed binary CS matrix (dashed lines), and this effect increases as the variance explained by the sigmoidal fits increases.

#### Known regulators of TF activity

The TFKO dataset contains 1,484 samples from strains in which a gene was deleted, including many known regulators of TF activity. Our next evaluation task was to determine the target TFs in samples where a known regulator of TF activity is perturbed. To evaluate performance on this task, we compiled a map of known TF activity regulators and their targets (File S31). For each TF, an activity regulator was assigned to it if there was published literature that proposed a direct interaction that affects TF activity by a specific mechanism, such as phosphorylation, nuclear localization, or complex formation. Networks, constraints, and initial fitting used the ZEV dataset. The resulting CS matrix was then used to fit TFAs and baselines to the 194 TFKO samples in which a known TFA regulator was perturbed. In each perturbation sample, all TFs were ranked by the absolute difference between their standardized log activities in the perturbed and unperturbed samples. The absolute value was used because the literature is not always clear on the direction of regulation. We then plotted the fraction of literature-supported targets that were ranked above a given percentile in the sample where their TFA regulator was perturbed (Fig. 4C). The optimized CS matrices (solid lines) identify more known TFA regulators than the signed binary matrices (dashed lines), especially at rank percentiles above 85%. The ChIP-PC network (blue) also outperforms the DE-PC (orange) by this metric.

#### ZEV time course data

Although we have focused on the ZEV data from 15 minutes after TF induction, they are part of time courses with samples taken 2.5, 5, 10, 15, 20, 30, 45, 60, and 90 minutes after induction (some timepoints are not available for some TFs). The expression data show that each TF is rapidly induced to a very high level and remains highly expressed throughout the 90 minutes. Therefore, we decided to infer TFAs for the entire time course, using a CS matrix derived from the TFKO data. For each time point, we computed a log fold change of the induced TF’s inferred activity, relative to its activity at time 0. We then fit a 4-parameter, sigmoidal, saturating curve to the time series (Methods; Fig. 4D). To eliminate curves that fit poorly, we tried several different thresholds on the variance explained by the sigmoidal fit. For each threshold, we calculated the fraction of fits that showed increasing activity throughout the time series (expected behavior) rather than decreasing activity. Overall, the TFAs with optimized CS matrices (solid lines) performed better than those with signed binary CS matrices (dashed lines) and this effect got stronger for curves with better fits (Fig. 4E). Furthermore, the DE-PC network (orange lines) performed substantially better than the ChIP-PC network (blue lines).

### Condition-independence of control strengths

Control strengths are intended to be condition-independent, quantitative measures of each TF’s potential to regulate each target gene. We have seen that good performance on TFA inference tasks can be obtained by using control strengths optimized on a different data set (Fig. 1A, 2A, 3, 4). Another indication that inferred CSs are transferable between data sets is that using the CS values inferred from one data set for TFA inference in a second data set increases the variance explained by 5%, relative to using the signed binary CSs (Fig. S2).

Another way to test the condition-independence of control strengths is to calculate the correlation between CSs inferred from two data sets collected in different growth conditions. To maximize the number of TFs and their target genes we could use, we first created a large network from the union of edges from ChIP-PC and the two DE-PC networks, one derived from the TFKO data and the other from the ZEV. After filtering out edges with conflicting sign constraints (see Methods) and dropping two TFs that were left with only a single target, this new network (Union-PC, File S25) contains 94 TFs, 1,416 target genes, and 2,731 edges. We optimized both TFA and CS matrices using both the TFKO and the ZEV datasets together (see Methods). As an initial validation of the resulting CS matrix, we used it to infer TFAs in a new data set consisting of 69 double-deletion strains (51) (See supplement). The results for predicting the direction of perturbation (86.4% correct) and identifying the perturbed TF (rank percentile 95.7%) were even better than the results for smaller networks (compare to Fig. 2A). The percentage of TFs that showed positive correlation between their mRNA levels and their inferred TFAs, 66%, is slightly below the results for the smaller networks, but still much better than chance. Readers can use the optimized CS matrix (File S26) for TFA inference in other datasets.

To calculate the correlation of CS values inferred in two growth conditions, we re-optimized Union-PC using only the TFKO data (synthetic complete medium with 2% glucose and no nutrient limitation) and then only the ZEV data (minimal medium with 2% glucose in phosphate-limited chemostats). Focusing on the 75 TFs with at least 5 targets and averaging across bootstrap samples of target genes for each TF, the median correlation was +0.32 and 61 of 75 TFs had positive correlations (81%). This shows that, while there may be some over-fitting of CSs to a particular growth condition, there is also a substantial amount of condition-independence.

### Evaluating control strengths directly

As another evaluation, we asked whether the control strengths inferred for the targets of a TF would correspond in any way to the strength with which the TF binds to the promoters of those targets in genome-wide binding location data. To do this, we turned to binding data obtained by the transposon calling cards method (52, 53). In this method, a TF is linked to a transposase, which deposits a transposon in the genome near where the TF is bound. The number of transposons in a gene’s promoter is an approximate measure of the amount of time the TF spends bound to that promoter. We predicted that promoter occupancy would be positively correlated with the inferred control strength, in most cases. Importantly, we considered only the genes that were targets of a TF in the input network and therefore had inferred control strengths. The input network itself does not contain quantitative binding strength information. We optimized both TFA and CS using the Union-PC network and both the TFKO and ZEV data together. Of the 11 TFs with Calling Cards data and at least 5 target genes, 9 (82%) have positive correlation between measured binding events and inferred CS value, with a median correlation of 0.31.

### Analyzing time courses after glucose influx

Next, we used the CS matrix for Union-PC (File S26) to infer baselines and TFAs for three expression time courses after yeast cells were provided with glucose. In the first dataset, yeast cells growing in galactose-limited chemostats were provided glucose to a final concentration of 0.02%(w/v) or 0.2% (54). In the second, batch cultures depleted glucose over a 24hr growth period before being transferred to fresh media with 2% glucose (55). We plotted the inferred activity of each TF as a function of time, fit both a 4-parameter sigmoid curve (as in Fig. 4) and a 6-parameter impulse curve (56), chose one of the two by the Bayes Information Criterion (57), and filtered out poor fits (R^2^<80%) (see Supplement). For four well-studied TFs, Gcr2, Gln3, Gcn4, and Msn2, the inferred TFA levels and fits for all 3 time courses are shown as points and lines inside turquoise circles in Figure 5. The shapes of the activity curves in response to glucose made sense: Gcr2, an activator of glycolytic genes, increased activity upon glucose addition; Gln3, Gcn4, and Msn2, activators of genes needed during nutrient deprivation, generally decreased activity upon glucose addition (Gln3 showed a very small increase in one time course). Gcr2 activity also makes sense in terms of the glucose concentrations added, returning to baseline quickly at the lowest concentration, more slowly at the intermediate concentration (note the last gold data point, which is not reflected in the curve), and not at all at the highest concentration.

**Figure 5.**
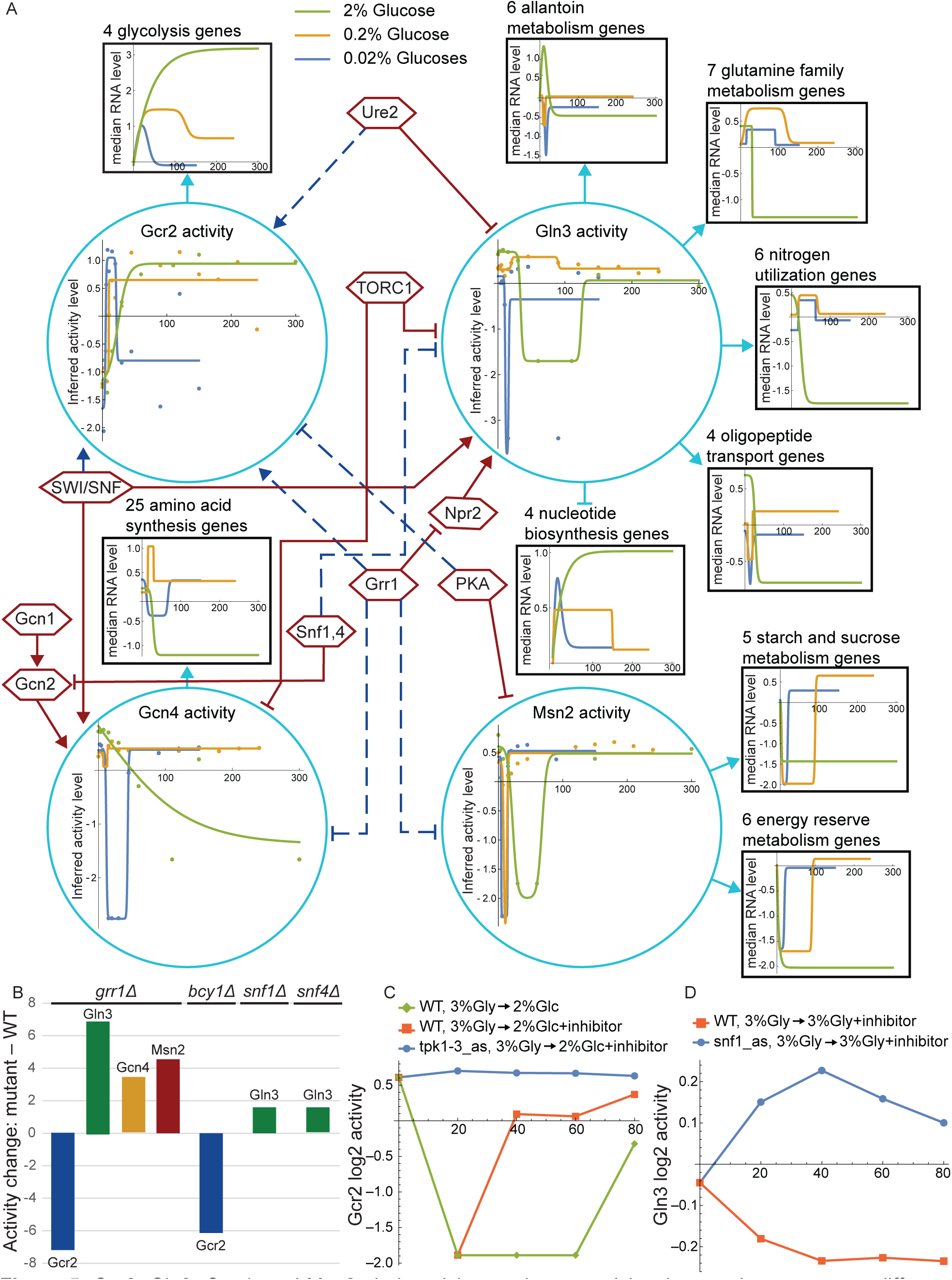
Gcr2, Gln3, Gcn4, and Msn2, their activity regulators, activity changes in response to different glucose concentrations, target genes sets, and target set expression patterns. A. Turquoise circles: Changes in inferred TF activity after addition of 2% glucose to post-diauxic-shift shake-flasks with synthetic complete medium (green) or addition of 0.2% (gold) or 0.02% (blue) glucose to cultures grown in galactose-limited chemostats with minimal medium. Points are Log2 of inferred activity level and lines are impulse or sigmoidal fits to the points, chosen by the Bayes Information Criterion. Black boxes: Sets of target genes that are regulated in the same direction and are annotated to a Gene Ontology or KEGG term enriched among targets of the TF that regulates them. Arrowheads indicate activation and T-heads repression. Colored lines are impulse or sigmoidal fits to the median Log2 fold-change of the annotated genes at each time point, relative to time 0. Hexagons: TF activity regulators inferred from analysis of two data sets as described in the text. Solid maroon lines indicate clear literature support while dashed blue lines indicate hypothesized novel edges. B. Change in activity of transcription factors in response to deletion of *GRR1, BCY1* (inhibitory subunit of PKA), *SNF1*, or *SNF4* (activating subunit of Snf1 complex). C. Gcr2 activity after addition of 2% glucose to cells growing on 3% glycerol. Glucose initially reduces Gcr2 activity, whether the cells are wild-type or mutants expressing analog-sensitive Tpk1-3. The shape of this response is different from Gcr2’s response to glucose under the conditions of Fig. 5A. Addition of the Tpk1-3 inhibitor with glucose to analog-sensitive cells eliminates that response, suggesting that PKA represses Gcr2 activity. This is consistent with the observation that deletion of *BCY1* reduces Gcr2 activity in 2% glucose (panel B). D. Gln3 activity is slightly elevated when inhibitor is added to cells growing in 3% glycerol and expressing an analog-sensitive Snf1, relative to WT cells, suggesting that the Snf1,4 complex represses Gln3 activity. This is consistent with the observation that deletion of either Snf1 or Snf4 increases Gln3 activity in 2% glucose (panel B).

To gain a deeper understanding of the network structure, the control strengths, and how they led to the TFA patterns shown in Figure 5, we carried out enrichment analysis on the network targets of the four TFs using Gene Ontology biological process annotations and Kyoto Encyclopedia of Genes and Genomes metabolic pathway annotations (58). We first discarded targets that, after optimization, had absolute CS <= 10^−4^, then analyzed the remaining activated and repressed targets of each TF separately, and removed redundant terms (59). All significantly enriched annotations that apply to four or more target genes are listed above the black squares in Figure 5. Each TF is connected to a target square by an arrow if the corresponding annotation was enriched among its activated targets and a T-head if the annotation was enriched among its repressed targets. Gcn4 and Msn2 had no repressed targets and the few repressed targets of Gcr2 had no enriched annotations that applied to four or more targets. The annotations and directions of regulation of target sets made sense in terms of the known function of each TF (60-64). To gain a deeper understanding of the network structure, the control strengths, and how they led to the TFA patterns shown in Figure 5, we carried out enrichment analysis on the network targets of the four TFs using Gene Ontology biological process annotations and Kyoto Encyclopedia of Genes and Genomes metabolic pathway annotations (58). We first discarded targets that, after optimization, had absolute CS <= 10^−4^, then analyzed the remaining activated and repressed targets of each TF separately, and removed redundant terms (59). All significantly enriched annotations that apply to four or more target genes are listed above the black squares in Figure 5. Each TF is connected to a target square by an arrow if the corresponding annotation was enriched among its activated targets and a T-head if the annotation was enriched among its repressed targets. Gcn4 and Msn2 had no repressed targets and the few repressed targets of Gcr2 had no enriched annotations that applied to four or more targets. The annotations and directions of regulation of target sets made sense in terms of the known function of each TF (60-64).

For each gene set in Figure 5, we calculated the median log fold change in each time course (not shown) and fit sigmoidal or impulse curves to them (shown in black boxes). In general, the shapes of the inferred activity curves for each TF made sense in light of the expression patterns of their target groups. For example, the inferred activity patterns of Gcr2 mirror those of the glycolysis genes it activates. Note that curves are expected to be inverted for repressed targets and that some of the lines occlude others at early time points, hiding early dips. Although the mRNA levels of target genes show a clear relationship to inferred activity levels, they do not always mirror each other perfectly. For example, the amino acid synthesis genes spike upward after addition of 0.2% glucose, whereas the inferred activity of Gcn4 dips and returns, presumably responding to influential target genes whose individual expression levels do not follow the pattern of the median levels shown.

Finally, we attempted to identify key regulators driving the TFA activity patterns observed in Figure 5, by inferring TFAs from two more data sets, using the same CS matrix. The first returns to the TFKO dataset for expression profiles of strains deleted for 567 different regulatory factors, including many known TF activity regulators (45). The second includes time courses after addition of rapamycin (an inhibitor of TORC1) or inhibitors of Snf1, Tpk1-3, or Sch9 (65). By examining the effects of these perturbations on the inferred activities of Gcr2, Gln3, Gcn4, and Msn2 and comparing them to published literature, we were able to confirm many previously described regulatory interactions and to identify a few potentially new ones (Fig. 5). We confirmed that Gcn2 activates Gcn4 (61-63, 66), Gcn1 activates Gcn4 (probably via its effects on Gcn2 (67)), and Ure2 represses Gln3 (probably by anchoring it to the plasma membrane, (62, 63)). We also saw evidence that Ure2 may activate Gcr2 directly or indirectly. We confirmed the well-known role of TORC1 as a repressor of Gln3 and Gcn4 activity (60, 64, 68) but did not see unequivocal evidence that it represses Msn2 as previously reported (60, 64). Our analyses showed that Grr1 represses Gln3, probably via Npr2 (69), which our analysis confirms as an activator of Gln3 (63)), and activates Gcr2 (probably indirectly; Fig. 5B). Although it has not been previously reported that Grr1 activates Gcr2, Grr1 is known to be required for glucose suppression, so activation of glycolytic genes via Gcr2 would be a consistent role. We also found that Grr1 represses Gcn4 and Msn2 (Fig. 5B), consistent with its being active in nutrient-replete conditions. Since Grr1 activates Gcr2 and represses the other three, a transient spike in its activity upon glucose influx could explain the upward and downward spikes we see in the activities of the four TFs. We confirmed that the SWI/SNF chromatin remodeling complex contributes to the activities of Gln3 (70, 71) and Gcn4 (61) and discovered that it also works with Gcr2. We confirmed that PKA represses Msn2 (60, 62-64, 72) and saw evidence that it also represses Gcr2 (Fig. 5C), which has not been previously reported. We confirmed that Snf1 represses Gcn4 in the absence of glucose and amino acids and saw evidence that it weakly represses Gln3 (Fig. 5B, D), contrary to previous claims that Snf1 activates Gln3 (64, 73). In summary, we have identified several likely regulators of Gcr2 activity, about which little was previously known, and discovered that Grr1, previously known for its role in glucose repression, is probably a positive regulator of glycolysis (via Gcr2) and a negative regulator of stress-induced TFs. These novel observations, which were mined from gene expression data via TF activity inference, constitute a rich trove of hypotheses for future experimental investigation.

## DISCUSSION

The ability to accurately infer changes in TF activity from changes in gene expression profiles provides a vital tool in the systems-biology toolbox. It enables us to look inside the cell, seeing not only the output of the circuits that control gene expression, but also their internal state. Observing the internal states of regulatory circuits is the key to understanding how these circuits control the cell’s transcriptional program in response to internal and external signals. We demonstrated how this tool can be used to gain new insights into the regulation of TF activity, such as the probable activation of Gcr2 and repression of Gcn4, Gln3, and Msn2 by the Grr1 ubiquitin ligase in the presence of glucose (Fig. 5). In future work, we hope to automate this process of identifying TFA regulators and develop genome-scale benchmarks with which to evaluate it.

Previous studies introduced various versions of the matrix factorization approach (1, 6, 8, 24). Here, we presented the first objective, genome-scale evaluation of this approach, using multiple measures of accuracy and multiple independent data sets. We found that, when inferring TFA and CS matrices, it is essential to have data in which TFs are directly perturbed and constraints derived from them. We also showed that a CS matrix derived from the TFKO and ZEV data can be used to successfully analyze other expression data sets that do not contain direct perturbations of TF activity. In two of our core metrics, matrix factorization with constraints yielded TFA values that cover more than half the distance from random (50%) to ceiling (100%; Fig. 2A). However, this probably underestimates the true accuracy of the method, since even perfect TFA inference would not necessarily yield 100% on these metrics. For example, deletion of an inactive TF will not result in decreased activity, and perturbation of a TF’s activity could lead to an equal or bigger change in the activity of a downstream transcription factor, pushing the true rank percentile of the perturbed TF below 100%. Similarly, the true percentage of TFs whose activity is positively correlated with their mRNA level is almost certainly less than 100%, due post-transcriptional regulation.

Building the input network from binding specificity models (e.g. PWMs) is a popular approach that is condition-independent in principle, but it did not do well in our evaluation (Fig. 2A). Network structures derived from either perturbation-response data (DE) or currently available binding location data (ChIP) perform about equally when using the highest-scoring edges, but as more edges are added, the DE network is more robust (Fig. 3). For the ChIP-PC network, we observed the best performance when using only the top 1250 edges, even though the recommended P-value threshold of 0.001 includes at least three times more. Using edges ranked 2001 through 3250 (Fig. 3A-C, Block 2) resulted in decreased performance on all metrics. The extended ChIP-PC networks of 77-80 TFs, which considered the top 2000 edges rather than the top 1250, also resulted in decreased performance on all the metrics (Fig. 3D). Since perturbation-response data are needed for sign constraints in any case, generating binding location data in addition may not be worth the effort and expense required. In other words, perturbation-response data is both necessary and sufficient for good performance.

Another recent paper (42) evaluated TFA inference algorithms by using expression data after TF knockdowns in human cell lines and *E. coli*, similar to our second metric. It reported that the perturbed TF was rarely among those with the greatest inferred activity changes and there was very little agreement among the algorithms tested. One factor that likely contributes to the difference in findings is that the algorithms they tested did not use sign constraints on controls strengths and could not distinguish between increasing and decreasing activity. In the absence of sign constraints, we also saw poor performance. Another difference is that the input networks they used were largely based on manual curation rather than automated processing of high-throughput data, so they lacked confidence scores, making it impossible to select only the most confident edges. We saw performance degrade when less confident edges were included in the network. Finally, their evaluation was carried out using perturbations to only a handful of TFs for evaluation, making the findings vulnerable to sampling error.

We were surprised to find that, by our three basic metrics, optimizing control strengths is not necessary for achieving good performance -- signed binary control strengths, taken directly from the input network and sign constraints, do almost as well (Fig. 4A,B). Optimized CS matrices result in somewhat better performance on two other metrics -- detecting literature-supported TFA regulators (Fig. 4C) and detecting the trend in TF activity from a time course (Fig. 4D,E). Furthermore, we found evidence that the optimized control strengths correlated positively with those learned by optimizing on data from different growth conditions and with the strength of binding to target promoters in independent binding data. Nonetheless, these correlations were far from perfect, so the limited impact of optimizing CS matrices on TFA accuracy may reflect the fact that the control strengths in our testing framework are optimized on one data set while TFAs are optimized and evaluated on another (Fig. 1). In many real applications, control strengths and activities would be optimized on the same data set. Improvements to the mathematical model could also increase the importance of CS optimization (see below). For now, however, using signed binary control strengths may be a reasonable choice for some organisms, especially when a limited amount of gene expression data is available. When control strengths are not optimized, the overall optimization changes from non-linear to linear, making it much faster and simpler.

Any approach to TFA inference relies on having a reasonably large and accurate set of targets for each TF. This can be challenging for TFs that are not very active in any of the conditions in which the data used to build the network were obtained. For example, the well-known TF Gal4 had only 3 targets in the Union-PC network. One of those, *GAL10*, is an established target with a known role in galactose metabolism, but CS optimization reduces the link between Gal4 and *GAL10*, emphasizing instead the link between Gal4 and dubious ORF YDR544C. This may be due to the fact that none of the samples to which we fit the activity levels were grown with galactose, so the expression of *GAL10* does not vary much. As a result, there is little need to explain *GAL10* expression as resulting from changes in Gal4 activity. A possible approach to this problem would be to discard inferences about TFs that have two or fewer significant targets after CS optimization.

Analysis of the error patterns on our three core benchmarks showed that poor performance was highly correlated with the fraction of a TF’s targets that exhibited opposite signs in the TFKO and ZEV data sets (Fig. 2B-D). Apparent sign conflict can occur when the true sign of regulation is consistent, but the effect of the perturbation on the target gene is so weak that random measurement noise leads to a sign error. This can be remedied by using multiple perturbation data sets to determine sign and discarding edges with sign conflicts, as we did when constructing the Union-PC network. However, conflicts can also occur because some TFs are repressors in some conditions and activators in others. For example, Rgt1 represses *HXT1* in low glucose but activates it in high glucose (74). This points to a limitation of any model that constrains TFAs to be non-negative and CSs to be one sign or the other. A possible solution would be to release the non-negativity constraint on TFs when there is sufficient evidence of a true sign change that applies to most of its targets.

The matrix factorization approach has several limitations. First, predicted gene expression does not saturate as TF activity gets large, whereas in reality each gene has maximum and minimum expression levels and the binding sites for each TF eventually become fully occupied. Second, the model assumes that each TF-target relationship is either activating or repressing in all conditions. Third, TF-TF interactions, such as competitive or cooperative binding, are not accounted for. Fourth, as parameters are optimized during the fitting process, the variance explained is not a good predictor of a model’s value for accurate TFA inference. Finally, when control strengths learned on one data set are used to model a different data set, the variance explained in the second data set is much lower than in the first. This suggests a degree of over fitting that might be remedied by parameter shrinkage. However, improving the variance explained in cross-validation is not guaranteed to improve the accuracy of the TFA parameters learned.

We found that TFA inference in yeast works reasonably well when best practices are followed, but there is still room for improvement. We anticipate improvements coming from better network maps. One likely source of better maps is new, more accurate methods for measuring TF binding locations (53, 75-78). The input network could also be improved by obtaining TF perturbation data from cells grown in new conditions. More improvement in TFA inference could come with better sign constraints and the possibility of allowing negative TF activity when the data strongly justify it. Mathematical models that more closely reflect the underlying biochemistry could also lead to better results, although such models come with new challenges. As new approaches are tried, the benchmarks presented here can be used to determine whether they robustly improve the accuracy of TFA inference, across multiple data sets and network maps.

## METHODS

### Model fitting

In initial fits (Fig. 1A, top), models are fit to gene expression data by least-squares linear regression, alternating between TFA and CS matrices (which include baselines), starting from 20 random initializations of the CS matrix. If a TF does not regulate a gene in the TF network map, the corresponding CS is held at zero; otherwise, it is constrained to be either positive (activating) or negative (repressing). If a TF is deleted in a sample, its activity is held at zero; if it is overexpressed, its activity is constrained to be greater than its activity in unperturbed samples. Except for deletion samples, all activities are constrained to be >= 0.0001. When learning a CS matrix, the mean activity of each TF, across all samples, is constrained to be one, since scaling a TF’s activities and control strengths by inverse factors does not affect the predictions. After each iteration, the non-baseline control strengths are held constant while the TFAs and baselines are fit to the second data set by alternating linear regression without constraining the mean activity of a TF (Fig. 1A, middle). Optimization against the first dataset is halted when R^2^ in the second data set peaks or after 100 iterations, whichever comes first. This halting criterion does not use the perturbation key of the second dataset. Comparable results were obtained by using Knitro (44), a general non-linear solver, to optimize all parameters at once.

### Ranking TFs

To calculate the rank percentile of a perturbed TF, the log2 of the activity values for each TF are standardized across all samples. In TF knockout samples, the standardized log2 activity values of each TF are sorted lowest (most negative) to highest (most positive); in TF overexpression samples, they are sorted from highest to lowest. The rank percentile is 100 - (rank - 1)/numTFs. The same procedure is used for identifying targets of TFA regulators (Fig4C), except that the sorted values are absolute difference of standardized log2 activity of each TF in each perturbed sample from the non-perturbed sample.

Code for this evaluation is available at https://github.com/BrentLab/TFA-evaluation

### Correlation between TFA and TF-mRNA

For each dataset, the second fittings (Fig. 1A, middle) included an unperturbed sample and 179 samples in which a TF was perturbed. For 1,000 bootstraps of the 180 samples, the fraction of TFs whose mRNA and TFA values were positively correlated was calculated, and the median fraction was reported. TFA values were not logged or standardized before calculating correlation. Code for this evaluation is available at https://github.com/BrentLab/TFA-evaluation

### Union-PC

Combined, the ChIP-PC network with sign constraints from either TFKO or ZEV and DE-PC with edges and sign constraints from either TFKO or ZEV contain 3,133 unique edges between 96 TFs and 1592 target genes. Edges with conflicting sign constraints from different networks were dropped and TFs with only one target gene were removed, along with their target gene, leaving 2,731 edges between 94 TFs and 1,416 target genes. When optimizing this network on both datasets together, the perturbation samples for each network TF in each dataset were included, along with an unperturbed sample. CS parameters were shared, but different baseline values were used for each dataset to compensate for potential constant shifts in gene expression measurements.

## Supporting information

S1_50-TF_ChIP-CC_binary_network_with_signs_from_TFKO

S2_50-TF_ChIP-CC_binary_network_with_signs_from_ZEV

S3_50-TF_ChIP-CC_network_with_optimized_values_from_TFKO

S4_50-TF_ChIP-CC_network_with_optimized_values_from_ZEV

S5_50-TF_TFA_values_optimized_with_ChIP-CC-TFKO_on_ZEV

S6_50-TF_TFA_values_optimized_with_ChIP-CC-ZEV_on_TFKO

S7_50-TF_ChIP-PC_binary_network_with_signs_from_TFKO

S8_50-TF_ChIP-PC_binary_network_with_signs_from_ZEV

S9_50-TF_ChIP-PC_network_with_optimized_values_from_TFKO

S10_50-TF_ChIP-PC_network_with_optimized_values_from_ZEV

S11_50-TF_TFA_values_optimized_with_ChIP-PC-TFKO_on_ZEV

S12_50-TF_TFA_values_optimized_with_ChIP-PC-ZEV_on_TFKO

S13_50-TF_DE-PC_binary_network_with_signs_from_TFKO

S14_50-TF_DE-PC_binary_network_with_signs_from_ZEV

S15_50-TF_DE-PC_network_with_optimized_values_from_TFKO

S16_50-TF_DE-PC_network_with_optimized_values_from_ZEV

S17_50-TF_TFA_values_optimized_with_DE-PC-TFKO_on_ZEV

S18_50-TF_TFA_values_optimized_with_DE-PC-ZEV_on_TFKO

S19_50-TF_PWM-PC_binary_network_with_signs_from_TFKO

S20_50-TF_PWM-PC_binary_network_with_signs_from_ZEV

S21_50-TF_PWM-PC_network_with_optimized_values_from_TFKO

S22_50-TF_PWM-PC_network_with_optimized_values_from_ZEV

S23_50-TF_TFA_values_optimized_with_PWM-PC-TFKO_on_ZEV

S24_50-TF_TFA_values_optimized_with_PWM-PC-ZEV_on_TFKO

S25_94-TF_Union-PC_binary_network_with_signs_from_TFKO_and_ZEV

S26_94-TF_Union-PC_network_with_optimized_values_from_TFKO_and_ZEV

S27_94-TF_TFA_values_optimized_with_Union-PC-TFKO-ZEV_on_2.0p_glucose

S28_94-TF_TFA_values_optimized_with_Union-PC-TFKO-ZEV_on_0.02-0.2p_glucose

S29_94-TF_TFA_values_optimized_with_Union-PC-TFKO-ZEV_on_Regulators

S30_94-TF_TFA_values_optimized_with_Union-PC-TFKO-ZEV_on_Zaman

S31_Literature_curated_regulators_of_TFA

## LIST OF ABBREVIATIONS

CS: Control Strength
TF: Transcription factor
TFA: Transcription factor activity
TFKO: gene expression dataset of deletion strains
ZEV: gene expression dataset of induced over-expression
ChIP-CC: network based on ChIP binding scores and TF-target mRNA correlation
ChIP-PC: network based on ChIP binding scores and TF perturbation response
DE-PC: network based on DE of target genes in response to TF perturbation
PWM-PC: network based on PWM motif scores and TF perturbation response

## DECLARATIONS

The authors declare no competing interests.

## ACKNOWLEDGMENTS

We thank Dhoha Abid for her help with data analysis, and we thank Artelys Knitro for their help in testing the optimization of our TFA inference model with their non-linear optimizer.

## AUTHORS’ CONTRIBUTIONS

CM and MB conceived of the research and wrote the paper. CM carried out all the coding and computational experiments.

## AUTHORS’ INFORMATION

CM is a computer science PhD student supervised by MB. MB is the Henry Edwin Sever Professor of Engineering at Washington University in Saint Louis and Professor of Genetics at the Washington University School of Medicine.

