## Supplemental Methods and Figures for "Inferring TF activities and activity regulators from gene expression data with constraints from TF perturbation data"

SUPPLEMENTARY FILES

S1_50-TF_ChIP-CC_binary_network_with_signs_from_TFKO.csv

S2_50-TF_ChIP-CC_binary_network_with_signs_from_ZEV.csv

Signed binary networks of edges between TFs and target genes. Edges come from top scores in ChIP dataset (1), while signs come from the correlation between TF-mRNA and target gene mRNA in either the TFKO (2) or the ZEV (3) dataset. See Network construction below for more details.

S3_50-TF_ChIP-CC_network_with_optimized_values_from_TFKO.csv

S4_50-TF_ChIP-CC_network_with_optimized_values_from_ZEV.csv

Optimized control strength values for edges between TFs and target genes. Edges and signs come from the binary networks (Files S1 and S2), while the quantitative values come from fitting to gene expression profiles of either the TFKO or the ZEV dataset.

S5_50-TF_TFA_values_optimized_with_ChIP-CC-TFKO_on_ZEV.csv

Optimized TFA values on ZEV dataset, using the ChIP-CC values that were optimized on TFKO.

S6_50-TF_TFA_values_optimized_with_ChIP-CC-ZEV_on_TFKO.csv

Optimized TFA values on TFKO dataset, using the ChIP-CC values that were optimized on ZEV.

S7_50-TF_ChIP-PC_binary_network_with_signs_from_TFKO.csv

S8_50-TF_ChIP-PC_binary_network_with_signs_from_ZEV.csv

S9_50-TF_ChIP-PC_network_with_optimized_values_from_TFKO.csv

S10_50-TF_ChIP-PC_network_with_optimized_values_from_ZEV.csv

S11_50-TF_TFA_values_optimized_with_ChIP-PC-TFKO_on_ZEV.csv

S12_50-TF_TFA_values_optimized_with_ChIP-PC-ZEV_on_TFKO.csv

See description for files S1-S6. Signs for the binary networks come from the gene expression response of targets to the perturbation of TFs in either the TFKO or the ZEV dataset. Specifically, if a target gene decreases or increases expression between a TF deletion sample and the WT, the edge between the TF and target gene is positive or negative respectively. If a target gene decreases or increases expression between a TF induced sample and the WT, the edge between the TF and target gene is negative or positive respectively.

S13_50-TF_DE-PC_binary_network_with_signs_from_TFKO.csv

S14_50-TF_DE-PC_binary_network_with_signs_from_ZEV.csv

S15_50-TF_DE-PC_network_with_optimized_values_from_TFKO.csv

S16_50-TF_DE-PC_network_with_optimized_values_from_ZEV.csv

S17_50-TF_TFA_values_optimized_with_DE-PC-TFKO_on_ZEV.csv

S18_50-TF_TFA_values_optimized_with_DE-PC-ZEV_on_TFKO.csv

See description for files S7-S12. Signs and edges for the binary networks come from the gene expression response of targets to the perturbation of TFs in either the TFKO or the ZEV dataset.

S19_50-TF_PWM-PC_binary_network_with_signs_from_TFKO.csv

S20_50-TF_PWM-PC_binary_network_with_signs_from_ZEV.csv

S21_50-TF_PWM-PC_network_with_optimized_values_from_TFKO.csv

S22_50-TF_PWM-PC_network_with_optimized_values_from_ZEV.csv

S23_50-TF_TFA_values_optimized_with_PWM-PC-TFKO_on_ZEV.csv

S24_50-TF_TFA_values_optimized_with_PWM-PC-ZEV_on_TFKO.csv

See description for files S7-S12. Edges for the binary networks come from top scores of motif scanning, and signs come from the gene expression response of targets to the perturbation of TFs in either the TFKO or the ZEV dataset.

S25_94-TF_Union-PC_binary_network_with_signs_from_TFKO_and_ZEV.csv

S26_94-TF_Union-PC_network_with_optimized_values_from_TFKO_and_ZEV.csv

S27_94-TF_TFA_values_optimized_with_Union-PC-TFKO-ZEV_on_2.0p_glucose.csv

S28_94-TF_TFA_values_optimized_with_Union-PC-TFKO-ZEV_on_0.02-0.2p_glucose.csv

S29_94-TF_TFA_values_optimized_with_Union-PC-TFKO-ZEV_on_Regulators.csv

S30_94-TF_TFA_values_optimized_with_Union-PC-TFKO-ZEV_on_Zaman.csv

See description for files S7-S18. Signs and edges for the binary network comes from non-conflicting edges of ChIP-PC from TFKO, ChIP-PC from ZEV, DE-PC from TFKO, and DE-PC from ZEV. Network edges are optimized on both TFKO and ZEV. TFA values used CS values optimized on TFKO and ZEV to fit glucose influx data, all TFKO to include samples beyond just TF deletion strains, and a dataset used in the paper to further analyze TF regulation (4).

S31_Literature_curated_regulators_of_TFA.csv

A curated map of TFA regulators. The first two columns are the systematic and common names of TFs. The next two columns are the systematic and common names of the TFA regulators. The fifth column indicates the direction of effect from the regulation, with -1 if the regulator decreases the activity of the TF, 1 if the regulator increases the activity of the TF, and 0 if the direction is unknown or not straightforward, i.e. part of a feedback loop. The next column is a brief description of the regulatory mechanism, and the last column records the PMID of the paper that describes the regulatory relationship.

METHODS

Datasets Used

*Yeast TFKO data* The microarray expression data of 1,484 single gene knockout strains (2) was downloaded from

<http://deleteome.holstegelab.nl/data/downloads/deleteome_all_mutants_controls.txt> A sample using expression level of 0 for all genes was assumed in order to stand in for WT.

*Yeast ZEV induction data* The microarray expression data of 199 single gene ZEV induction strains (3) was downloaded from <https://storage.googleapis.com/calico-website-pin-public-bucket/datasets/pin_tall_expression_data.zip> Only the column labeled log2_cleaned_ratio was considered for this work. A sample using expression level of 0 for all genes was assumed in order to stand in for the 0min timepoint, when induction of over-expression had not yet started.

*Yeast ChIP-chip data* P-values that represent TF binding significance from ChIP-chip experiments (1) were downloaded from

<http://younglab.wi.mit.edu/regulatory_code/GWLD.html> Values were transformed to negative log10 p-values for the purpose of treating them as confidence scores, where greater values indicate greater support.

*Yeast PWM data* Position weight matrices for S. cerevisiae motifs in the ScerTF database (5) were downloaded from <http://stormo.wustl.edu/ScerTF/> Values from FIMO scanning (6) for motif hits were transformed to negative log10 p-values for the purpose of treating them as confidence scores, where greater values indicate greater support. If multiple hits were found between linking the same TF and target gene, the maximum score was used.

*Yeast double-deletions data* The microarray expression dataset of 69 double-deletion strains (7) was downloaded from <http://www.holstegelab.nl/publications/GSTF_geneticinteractions/> A sample using expression level of 0 for all genes was assumed in order to stand in for WT.

*Yeast time course data*

The microarray expression data of 9 time-points (0min, 3, 7.5, 15, 30, 60, 110, 150, 300min) after 2% glucose influx for WT strains (8) was downloaded from <http://www.holstegelab.nl/publications/glucose_regulatory_system/> A sample using expression level of 0 for all genes was assumed in order to stand in for 0min.

The microarray expression data of 13 (0min, 2, 4, 6, 8, 10, 15, 20, 30, 45, 90, 120, 150) and 15 (0min, 3, 5, 7, 10, 15, 20, 30, 45, 90, 120, 150, 180, 210, 240) time-points after 0.02% and 0.2% glucose influx for WT strains (9) was downloaded from GEO with accession ID GSE4158. A sample using expression level of 0 for all genes was assumed in order to stand in for 0min.

The microarray expression datasets of 5 time-points (0min, 20, 40, 60, 80) for multiple conditions and multiple strains (4) were downloaded from <https://puma.princeton.edu/cgi-bin/publication/viewPublication.pl?pub_no=524>. Where possible, all samples were re-scaled to use the WT in 3% glycerol condition as the reference, and a sample using expression level of 0 for all genes was assumed in order to stand in for this reference.

Network construction

All possible TF-target interactions are first ranked according to the strength of evidence that the TF regulates the potential target gene. We did this for yeast (*S. cerevisiae*), ranking edges according to their negative log p-value in a comprehensive ChIP-chip dataset, their absolute differential expression in a TF perturbation sample, or their maximum negative log p-value in a comprehensive PWM dataset. To integrate data sources, the edges can be rank-averaged at this point, though the performance of such integrated networks is not shown in this paper.

To build a network map, we first dropped all but the top 1,250 edges. Then starting from the top, edges are added until 50 TFs were included. If there are not enough TFs, the total number of edges considered are iteratively increased by 25 until at least 50 TFs can be recovered. Any remaining edges are only included if they emanate from TFs already in the map. This initial map is then checked for any TFs with a single target gene, and any set of TFs with identical target genes, to be removed, both TFs and target genes. The former was removed to avoid having TFA values dependent on only one feature, which would make it extremely vulnerable to noise or measurement error, while the latter was removed as identical target genes sets means the influence of the TFs are impossible to separate. If necessary, we then return to the list and add edges that were previously skipped over, repeating all steps until the network holds steady at 50 TFs.

The ChIP network considered the top 1,250 edges, using 1,104 of them to build a network of 50 TFs and 778 genes. The minimum score (negative log adjusted p-value) was 4.37.

For the DE network based on TFKO, the top 1250 edges were not sufficient to build a network of 50 TFs when using the TFKO dataset, so additional edges were considered in increments of 25 until the top 1400 edges was found to be sufficient. 1,283 of them were used to build a network of 50 TFs and 573 genes. The minimum score (absolute logFC of gene expression) was 1.25.

The DE network based on ZEV considered the top 1,250 edges, using 890 of them to build a network of 50 TFs and 686 genes. The minimum score (absolute logFC of gene expression) was 1.67.

The PWM network considered the top 1,250 edges, using 1,023 of them to build a network of 50 TFs and 894 genes. The minimum score (maximum negative log p-value) was 9.19.

To add correlation-based sign constraints (eg. ChIP-CC), the direction of correlation between the TF and target gene’s expression levels across samples was calculated. Samples where a network TF was directly perturbed were not included in the correlation calculation to ensure that these constraints were based on general correlation trends and not based on gene expression response to direct perturbation of an assigned TF regulator.

To add perturbation-based sign constraints (eg. ChIP-PC), the direction of a gene’s expression in the perturbation sample of its TF was used. For TFKO-based constraints, the sign was reversed to indicate that the TF likely activates a gene that decreases expression in its absence, and vice versa. For ZEV-based constraints, the sign was used directly.

For networks using lower quality edges, the goal was to build networks where the support for all edges decreased, without overlap. In anticipation of situations like the DE network based on ZEV, where the total number of edges had to be increased beyond 1250 in order to obtain a network of 50 TFs, a generous 2,000 edges were selected to comprise a “block,” even though the networks themselves never needed all 2,000 edges. Block 1 networks are the same networks described above, while the Block 2 networks were created after reassigning ranks when the top 2,000 edges were zero-ed out, the Block 4 networks were created after reassigning ranks when the top 6,000 edges were zero-ed out, etc... The number of edges considered to build each network was kept to 1,250 whenever possible, with the only exceptions being two DE networks based on TFKO, where Block 1 used 1,400 as described above, and Block 2 used 1,450 edges.

The ChIP and DE extended networks were built for direct comparison with combining the TFA values inferred from ChIP-PC and DE-PC. The union of ChIP-PC and DE-PC based on TFKO was 77 TFs, and the union of ChIP-PC and DE-PC based on ZEV was 80 TFs. Therefore, the ChIP and DE extended networks needed to cover the same number of TFs, depending on which dataset was being used for CS optimization instead of TFA validation. The number of edges considered was extended proportionally; eg. a network increase from 50 to 77 TFs started with an increase from 1,250 to 1250*(77/50) =1,925 edges.

The ChIP extended network for optimizing on TFKO considered the top 1,925 edges, using 1,866 of them to build a network of 77 TFs and 1,196 genes. The minimum score (negative log adjusted p-value) was 3.70.

The ChIP extended network for optimizing on ZEV considered the top 2,000 edges, using 1,968 of them to build a network of 80 TFs and 1,250 genes. The minimum score (negative log adjusted p-value) was 3.64.

The DE extended network for optimizing on TFKO considered the top 3,575 edges, using 3,284 of them to build a network of 77 TFs and 1,213 genes. The minimum score (absolute logFC of gene expression) was 0.85.

The DE extended network for optimizing on ZEV considered the top 2,000 edges, using 1,707 of them to build a network of 80 TFs and 1,019 genes. The minimum score (absolute logFC of gene expression) was 1.45.

The Union-PC network was built from the union of ChIP-PC edges, and edges from the two DE-PC networks, one derived from the TFKO data and the other from the ZEV. This started out as a set of 3,133 unique edges between 96 TFs and 1592 target genes. Among the edges in the ChIP network, 413 have conflicting sign constraints between the ZEV and TFKO datasets and were therefore discarded. Among the 42 edges in both the ZEV- and TFKO-based DE networks, none have conflicting sign constraints, so all edges were kept. Finally, among the 90 edges found in both the trimmed ChIP network and DE networks, none had conflicting sign constraints, which left us with 2,732 edges. After filtering out two TFs that were left with only a single target gene due to the loss of ChIP edges, we were left with 94 TFs, 1,416 target genes, and 2,731 edges.

Model-Fitting

Given a network of edges between TFs and genes, random values between -10 and 10 are generated to stand in for the control strengths of those edges, as well as the baseline expression level of genes. Any sign constraints on control strength parameters are then checked, flipping the signs of the random control strength values as necessary. For this paper, this process was repeated to create 20 random starts for each network structure.

Each random start is used to create a linear optimization problem, where activity values are optimizable, non-negative parameters to minimize the squared error between measured gene expression and model predicted gene expression. For this paper, the gene expression set used in the initial fitting was composed of one perturbation sample for each of the network TFs and a WT sample. The activity of TFs deleted in a sample are set to zero, while the activity of TFs over-expressed in a sample are constrained to be greater than its activity in wild-type samples. In this model, multiplying a TF’s activity values by a scaling factor and dividing its control strengths by the same factor leaves the predicted expression levels unchanged. Since the scale cannot be determined from the expression data, we constrain the mean activity of each TF, across all samples, to be one. After optimization, the random control strength values are discarded, and a new linear optimization problem is created, where control strengths and baseline values are optimizable parameters to minimize the squared prediction error. The control strengths of TFs are constrained to be negative or positive if the edge is known to be activating or repressive, respectively. This back-and-forth process, known as an iteration of bi-linear optimization, is repeated until the variance explained from fitting the new control strength values on the second dataset peaks, or until a maximum of 100 iterations is reached.

Finally, across the random starts, the set of parameters that achieved the greatest variance explained on the optimization set is selected for evaluation.

When optimizing for new baseline and activity parameters using an existing set of control strength parameters, the bi-linear framework is maintained. New activity parameters are optimized first, and the mean activity of each TF across samples is not constrained to one. After optimizing for activity, new baseline parameters are optimized. This iterates until the improvement in variance explained falls below 1%. For this paper, the gene expression set for this second fitting consisted of 179 perturbation samples and a WT sample.

When optimizing the Union-PC network on both the ZEV and TFKO datasets, the bi-linear approach remains the same, where TFA values for all samples are optimized against 20 sets of random CS values, before new CS values are optimized against the optimized TFA values. However, due to the datasets coming from different conditions, labs, and microarray technology, different baselines were optimized for the different datasets to compensate for any constant shifts in gene expression measurements. One perturbation sample for each TF in the network from each of the two datasets was included in the gene expression set, plus a WT sample for each of the two datasets. The set of TFA and CS values that achieved the greatest variance explained on the combined ZEV and TFKO dataset was selected for evaluation.

All optimization in this paper was done using Gurobi, an optimizer that offers free Academic licenses (10).

Calculating Evaluation Metrics

To ensure the inferred TFA values being evaluated is entirely blind of the criteria being evaluated, optimization of model parameters is done in two steps. The initial step optimizes all parameters to fit a dataset, either TFKO or ZEV. During this optimization, constraints can be applied to the activity parameters based on the known perturbations in the optimization dataset, as well as to the control strength parameters based on the known DE of target genes.

For the second step, optimized control strength parameters are used to optimize new baseline and activity parameters for a separate dataset. This second optimization does not allow activity parameters to be constrained based on known perturbations in the new dataset, nor for the control strength parameters to be updated based on conflicting DE of target genes.

The first three metrics explained below can be evaluated using code available here:

https://github.com/BrentLab/TFA-evaluation

*Direction of Perturbation* To predict the direction of TF perturbation in a given sample, we compare the TF’s activity in that sample to its activity in the WT sample. The percent of samples correctly predicted is calculated for each train-test direction, with the final score as an average between the two. To calculate the p-value, a binomial test is calculated for a 50% random chance of guessing the correct direction, where the number of trials is the total number of TFs evaluated in each dataset, averaged using Fisher’s combined probability test.

*Median Rank Percentile* To predict the perturbed TF, we log and standardize all activities to Z-scores within each TF, then rank all standardized log activities in each sample. If a sample involves a TF overexpression, we rank from highest to lowest; if a sample involves a TF knockout or knockdown, we rank from lowest to highest. The rank 1 TF is given the rank percentile of 100%, while subsequent TFs are $\left( 100-\frac{\left( rank-1 \right)}{numTFs} \right)\%$. The rank percentile of the perturbed TF is used as an accuracy score for each sample, and the median rank percentile is used as an accuracy score for each dataset (TFKO and ZEV), with the final score being an average between the two train-test directions. To calculate the P-value, a binomial test is calculated where the number of trials is the number of samples evaluated, half are successes, and the probability of success is the probability of randomly achieving the final score or better as a rank percentile. This represents the desired null model of how likely it is that half the TFs achieve at least the median rank percentile, given the random probability of that rank percentile.

*Positive correlation* The fraction of TFs whose measured mRNA level and inferred activity are positively correlated is calculated for 1,000 bootstraps from the 180 samples used in the second fitting, and the median from the bootstrapping for each train-test direction is then averaged to return as the final score. To calculate the P-value, a binomial test is calculated for a 50% random chance of getting a positive correlation, where the number of trials is the total number of TFs evaluated in each dataset, averaged using Fisher’s combined probability test. The statistical significance of the individual TFs’ correlations is not considered, since the bootstrapping provides robustness against sampling error.

*Known regulators of TF activity* We compiled a map of TF activity regulators by curating literature on all network TFs, looking for published work that proposed a specific mechanism of activity regulation, such as phosphorylation, nuclear localization, or complex formation (File S31). To evaluate these literatures supported relationships, we rank the change in all standardized activities in samples where a known regulator of TF activity is perturbed compared to the wild-type sample. As the literature does not always give a clear, unique direction to the expected change in TF activity, we rank from highest to lowest absolute value. The rank percentile of the regulated TF is used as an accuracy score for the relationship, and the median rank percentile is used as an accuracy score for the whole dataset.

*ZEV time course data* The expected behavior of a TF in the yeast ZEV induction dataset is an increasing sigmoid in response to the induced transcription of its gene. To evaluate whether we can recover that expected response pattern, we inferred TFAs for the whole ZEV time course, using a network, constraints, and CS matrix derived from the TFKO data. For each time point, (2.5, 5, 10, 15, 20, 30, 45, 60, and 90 minutes after induction of TF overexpression, though not every time point was available for every TF), we computed a log fold change of the induced TF’s inferred activity, relative to the inferred activity at time 0. We then fit a 4-parameter, sigmoidal, saturating curve to the time series:

$$h_{0}+\left( \frac{h_{1}-h_{0}}{1+e^{\beta(x-t)}} \right)$$

To eliminate curves that fit poorly, we tried several different thresholds on the variance explained by the sigmoidal fit. For each threshold, we calculated the fraction of fits that showed increasing activity throughout the time series (expected behavior) rather than decreasing activity.

*Multiple Perturbed TF Identification task* For the double deletion set, we want to score our ability to predict the perturbed TFs in any sample where a network TF is deleted, so we first check each sample if the knocked-out TFs are in the network. If none are, we skip the sample, and if one is, we rank and score in the same way we would rank and score a sample with a single TF knock-out. If both TFs are in the network, the standardized log2 activity values are ranked twice, once without including the first TF, and once without including the second TF. The rank percentile of the perturbed TF is used as an accuracy score for each ranking, and the median rank percentile is used as the summary accuracy score.

*Calculating CS Condition Independence* To evaluate the condition independence of CS values, we wanted to calculate the correlation of CS values inferred in two different growth conditions. The Union-PC network was optimized using only the TFKO data (synthetic complete medium with 2% glucose and no nutrient limitation) and then only the ZEV data (minimal medium with 2% glucose in phosphate-limited, continuous-flow chemostats). Of the 94 TFs in the network, 75 of them had at least 5 targets, which we chose as a minimum since correlations across fewer data points would not be very meaningful. The average correlation for 1,000 bootstrapped samples of the target gene set is calculated for each TF, between the CS values optimized on the two datasets.

*Calculating CS correlation with binding data* To evaluate whether the control strengths inferred for the targets of a TF would correspond to the strength with which the TF binds those targets, we turned to binding data obtained by the transposon calling cards method (11, 12). In this method, a TF is linked to a transposase, which deposits a transposon in the genome near where the TF is bound. The number of transposons in a gene’s promoter is an approximate measure of the amount of time the TF spends bound to that promoter, which we assume to be correlated with true control strength, in most cases. The Union-PC network was optimized on both the TFKO and ZEV data together, and 11 of the 94 TFs have both Calling Cards data and at least 5 target genes. Using 1,000 bootstrap samplings of target genes, the median correlation between inferred CS and measured transposons was calculated for each TF.

Exploratory Analysis of Inferred TFA values

*Glucose influx response pattern* After inferring TFA values for three time-courses of glucose influx, we want to summarize the general behavior pattern of TFA response in each. Thus, we plot the inferred activity of each TF as a function of time, fitting both a 4-parameter sigmoid curve and a 6-parameter impulse curve (13), choosing one of the two by the Bayes Information Criterion (14). The sigmoid curve is the same as used for ZEV time-course response pattern analysis, while the impulse curve allows for a return to a new baseline level:

$$\frac{1}{h_{1}}\left( h_{0}+\frac{h_{1}-h_{0}}{1+e^{\beta\left( x-t_{1} \right)}} \right)\left( h_{2}+\frac{h_{1}-h_{2}}{1+e^{-\beta\left( x-t_{2} \right)}} \right)$$

This results in five general categories of behavior: an upward spike, a downward spike, monotonically increasing, monotonically decreasing, and poor fits (R^2^<80%). Using these categories, we could check for expected behavior in response to glucose, as well as consistent behavior across the time-courses.

*Response to perturbation of protein complexes* The regulatory relationship of complexes and TFs was partially explored using the TFKO dataset, which included knock-out perturbations of many components of complexes, like SWI/SNF and TORC1. In order to analyze the direction and magnitude of effect from perturbing the complex, the median change in standardized log2 activity for a TF across all samples where a component of the complex was knocked-out was compared to other TFs, and to the median change from random sets of the same number of samples.

Supplementary figures

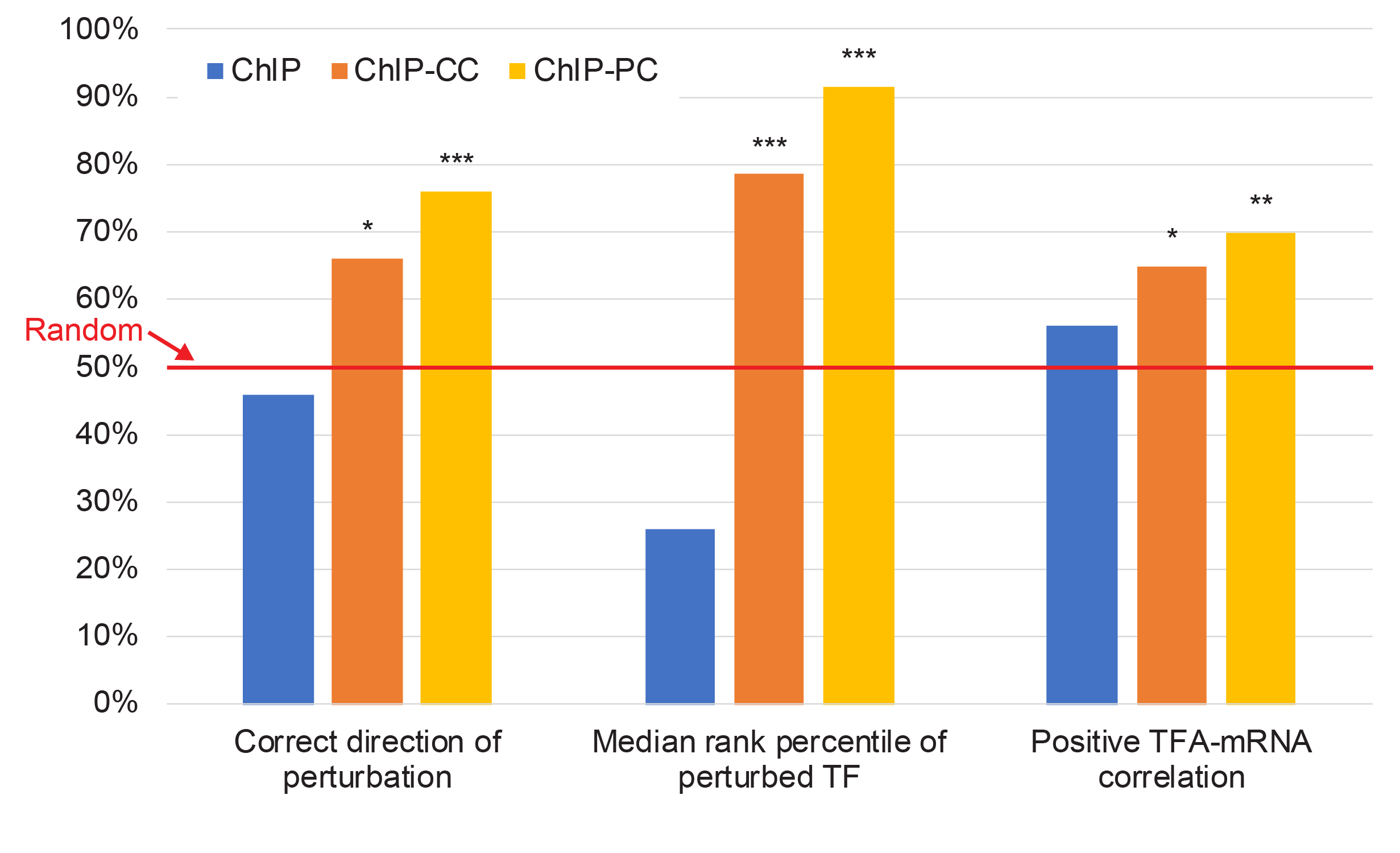

Supplementary Figure 1

Using ChIP to define the network without constraining the signs (blue) resulted in performance not significantly better than random. Performance of the ChIP network with correlation-based (orange) and perturbation-based (yellow) sign constraints are plotted for comparison.

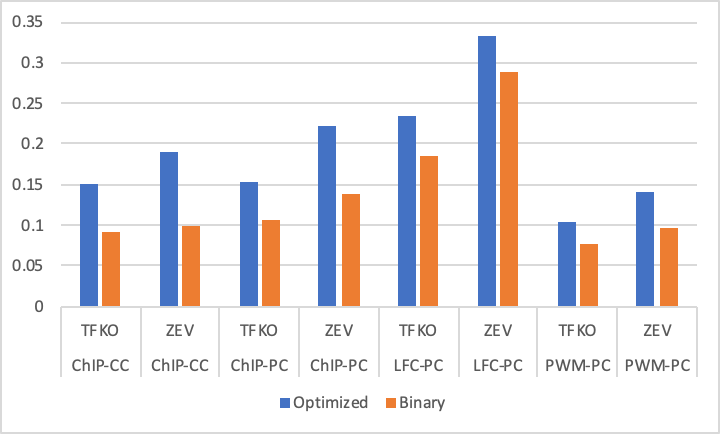

Supplementary Figure 2

Comparison of variance explained when using CS values optimized on a different data set (blue bars) or signed binary CS values (orange bars). Annotation below each pair of bars indicates the network and constraints used and the data set on which they were optimized. In all cases, both CS matrices are used to infer TFAs and baselines on the other data set and the variance explained is plotted. All pairs of bars show that CS matrices optimized on a different data set yield better fits than signed binary matrices, indicating that optimized CS matrices are, to some degree, transferrable from one growth condition to another.

10.	L. Gurobi Optimization, Gurobi Optimizer Reference Manual. (2020).

11.	H. Wang, D. Mayhew, X. Chen, M. Johnston, R. D. Mitra, Calling Cards enable multiplexed identification of the genomic targets of DNA-binding proteins. *Genome Res* **21**, 748-755 (2011).

12.	D. Mayhew, R. D. Mitra, Transposon Calling Cards. *Cold Spring Harb Protoc* **2016**, pdb top077776 (2016).

13.	G. Chechik, D. Koller, Timing of gene expression responses to environmental changes. *J Comput Biol* **16**, 279-290 (2009).

14.	G. Schwarz, Estimating the Dimension of a Model. *Ann. Statist.* **6**, 461-464 (1978).
